# DILP8 serves as a mature follicle sensor to prevent excessive accumulation of mature follicles in *Drosophila* ovaries and oocyte aging

**DOI:** 10.1101/2025.04.02.646814

**Authors:** Rebecca Oramas, Katarina Yacuk, Stella E. Cho, Natalie R. Aloisio, Jianjun Sun

**Affiliations:** Department of Physiology & Neurobiology, University of Connecticut, Storrs, Connecticut 06269 USA; Institute for Systems Genomics, University of Connecticut, Storrs, Connecticut 06269 USA

**Keywords:** mature follicle sensor, *dilp8*, *lgr3*, virgin ovulation, oocyte aging

## Abstract

Excessive mature follicle accumulation in ovaries harms oocyte health and offspring viability. As such, the number of mature follicles in ovaries is tightly controlled. In *Drosophila*, each ovary is comprised of ∼16 ovarioles, each containing 1-2 mature follicles, regardless of the female’s mating status. The mechanism by which the female flies count the number of mature follicles to coordinate egg release and oogenesis remains a mystery. Previous work, along with our RNAseq and antibody analysis, demonstrated that *Drosophila* insulin-like peptide 8 (DILP8) is expressed in somatic follicle cells of mature follicles but not in younger follicles. Contrary to previous findings, we found that DILP8 is not essential for mating-induced ovulation/egg laying. In contrast, global depletion or follicle-cell-specific knockdown of *dilp8* leads to defective ovulation/egg laying and a significant increase of mature follicles in virgin females. In addition, we found that excessive accumulation of mature follicles in *dilp8*-knockdown females leads to poor oocyte quality. Furthermore, knockdown of *Lgr3* (*Leucine-rich repeat-containing G protein-coupled receptor 3*), encoding a previously identified DILP8 receptor, showed similar ovulation/egg laying defects, accumulation of mature follicles, and poor oocyte quality in virgin females. Therefore, DILP8 functions as a mature follicle sensor to prevent excessive accumulation of mature follicles and maintain oocyte quality through the Lgr3 receptor in virgin females. Because mating is not always available in the wild and the DILP8/Lgr3 pathway is highly conserved across multiple species, our findings suggest that DILP8/Lgr3 is likely critical for maintaining the optimal reproductive fitness of virgin females and for species survival in the wild.

## Introduction

Oogenesis is a conserved reproductive process by which a germ cell develops into a mature, fertilizable oocyte (Spradling et al., 2022), which can be released from its surrounding somatic follicle cells through ovulation. In insects, this ovulation event is induced by mating so that the ovulated oocytes can be fertilized by sperm to maximize progeny production. Therefore, virgin females typically do not ovulate or ovulate less frequently than mated females to preserve mature follicles and prevent energy waste. However, prolonged preservation of mature follicles inside the ovary leads to oocyte aging and deceased offspring hatching rates (Greenblatt et al., 2019). The mechanisms by which females coordinate oogenesis and ovulation to ensure optimal accumulation of mature follicles in the ovaries while preventing oocyte aging and maximizing reproductive outcomes remains largely unknown. In other words, it is unclear how females sense their mature follicles. This question is especially significant given that mating opportunities in the wild are not always available despite insects utilizing multiple visual, audial, and chemical cues to find their mates.

*Drosophila* oogenesis has been well characterized in the past several decades (Berg et al., 2023). Each ovary is composed of ∼16 ovarioles, which are egg chamber (follicle) assembly lines with germline and follicle stem cells residing in the germarium at the anterior end and the most mature stage-14 follicles at the posterior end (Spradling, 1993). Progenies of germline and follicle stem cells in the germarium proliferate and differentiate to form a stage-1 egg chamber, which develops through 14 morphologically distinct stages to reach maturity and gain ovulatory competency. Our recent work demonstrated a transcriptional network including Ftz-f1, Single-minded (Sim), and Hindsight (Hnt) that promotes follicle maturation and ovulatory competency (Deady et al., 2017; Knapp et al., 2020; Oramas et al., 2023). Sim functions as a master regulator to turn on all ovulatory genes in follicle cells at stage 14 (Oramas et al., 2023), including the Octopamine receptor in mushroom body (Oamb), matrix metalloproteinase 2 (Mmp2), and NADPH oxidase (Nox). Mating induces octopamine (OA) release from the nerve terminal innervating the ovaries (Monastirioti, 2003; Heifetz et al., 2014; White et al., 2021), which activates Oamb in mature follicle cells and the oviduct to induce follicle rupture/ovulation (Deady and Sun, 2015; Lee et al., 2003; Lee et al., 2009). A wide array of work has also been done to illustrate the mechanism behind mating-induced behavior changes in *Drosophila* females. These include not only increased ovulation/egg laying but also reduced mating receptivity, changed feeding behavior, gut morphology, germline stem cell divisions, etc.--collectively referred to as the post-mating response (Avila et al., 2011; White et al., 2021). The post-mating response is mainly influenced by sex peptide (SP), a seminal fluid component transferred into the female reproductive tract after mating and acting on the SP receptor in the sensory neurons innervating the female reproductive tract (Liu and Kubli, 2003; Yapici et al., 2008; Hasemeyer et al., 2009; Yang et al., 2009; Wang et al., 2020). However, we do know that virgin female flies also ovulate, although less frequently. It is unclear what signal induces virgin ovulation and how virgin females control the number of mature follicle accumulation in their ovaries.

DILP8 is one of the eight insulin-like peptides found in *Drosophila* and has been extensively studied for its role in coordinating growth between organs and developmental timing (Gontijo and Garelli, 2018; Juarez-Carreño et al., 2018). DILP8 is highly secreted from larval imaginal discs with growth perturbation and mediates the delay of pupariation to ensure normal adult organ size (Colombani et al., 2012; Garelli et al., 2012; Hariharan, 2012). Depending on the nature of the perturbation, DILP8 expression is regulated by multiple signaling pathways, including JAK/STAT, Hippo, JNK, and Xrp1 (Boone et al., 2016; Boulan et al., 2019; Demay et al., 2014; Katsuyama et al., 2015; Sanchez et al., 2019). DILP8 mediates its functions by binding to the Leucine-rich repeat-containing G protein-coupled receptor 3 (Lgr3), a member of the highly conserved family of relaxin receptors. Lgr3 is expressed in two neurons in each brain lobe that innervate prothoracic triggering hormone-producing neurons, influencing prothoracic gland ecdysone synthesis and secretion. Additionally, Lgr3 functions directly in the prothoracic gland to regulate organ growth and developmental timing (Jaszczak et al., 2015; Jaszczak et al., 2016; Karanja et al., 2022). In normal development without perturbation, DILP8 is sharply induced at the larval-to-pupal transition by ecdysone signaling and acts on Lgr3+ neurons to ensure organ size adjustment and control the pupariation motor program for proper pupal formation (Blanco-Obregon et al., 2022; Heredia et al., 2021). In contrast to their role in development, DILP8 and Lgr3 are also expressed in adults, and their function in adult physiology is limited to a few studies. Tumor-derived DILP8 acts on Lgr3+ neurons in the brain to induce cancer anorexia, which is highly conserved in mammals (Yeom et al., 2021). Mating increases sugar intake by females via activating sexually dimorphic Lgr3+ neurons (Laturney et al., 2023; Meissner et al., 2016). In addition, DILP8 is specifically expressed in mature follicle cells of *Drosophila* ovaries (Li et al., 2023; Liao and Nässel, 2020; Tootle et al., 2011). Mutation of *dilp8* leads to a reduced number of eggs laid after mating, likely due to ovulation defects (Li et al., 2023; Liao and Nässel, 2020).

In this work, we characterized the role of ovarian DILP8. We found that DILP8 is a downstream target of Sim in mature follicle cells. Contrary to previous findings, we found that DILP8 in adult females is not essential for mating-induced ovulation and egg laying. Instead, ovarian DILP8 plays a critical role in promoting virgin ovulation, likely through the Lgr3 receptor. Ovarian DILP8 functions as a mature follicle sensor to prevent excessive accumulation of mature follicles and oocyte aging in virgin females.

## Results

### DILP8 is a downstream target of Sim in mature follicle cells

Our previous work showed that Sim is a master regulator for promoting follicle maturation and ovulation (Oramas et al., 2023). To identify more downstream targets of Sim in mature follicles, we carried out an RNAseq analysis and identified *dilp8*, which was significantly downregulated in mature follicles after *sim* knockdown (Figure S1A). We applied DILP8 antibody staining to further characterize DILP8 expression in the late oogenesis. DILP8 staining was barely detected in follicle cells of early stage-14 egg chambers, was significantly upregulated at mid stage 14, and reached the highest level at late stage 14 (Figure 1A-C). DILP8 staining was not detected in follicles before stage 14. These results are consistent with previous reports (Liao and Nässel, 2020; Tootle et al., 2011). We also confirmed the specificity of the DILP8 antibody by examining the DILP8 expression in follicle cells with *dilp8* knockdown via RNA interference (RNAi). Flip-out clones with *dilp8^RNAi^* expression showed low/no signal, while adjacent wild-type follicle cells showed high DILP8 expression at late stage 14 (Figure S1B-C). In addition, follicles with *dilp8^RNAi^*expression driven by *Vm26Aa-Gal4* in follicle cells from stages 10 to 14 showed no upregulation of DILP8 expression (Figure 1D-G, S1D). All these data suggest that DILP8 is upregulated in stage-14 (mostly mid and late stage-14; mature follicles) and can be efficiently knocked down using RNAi.

**Figure 1.**
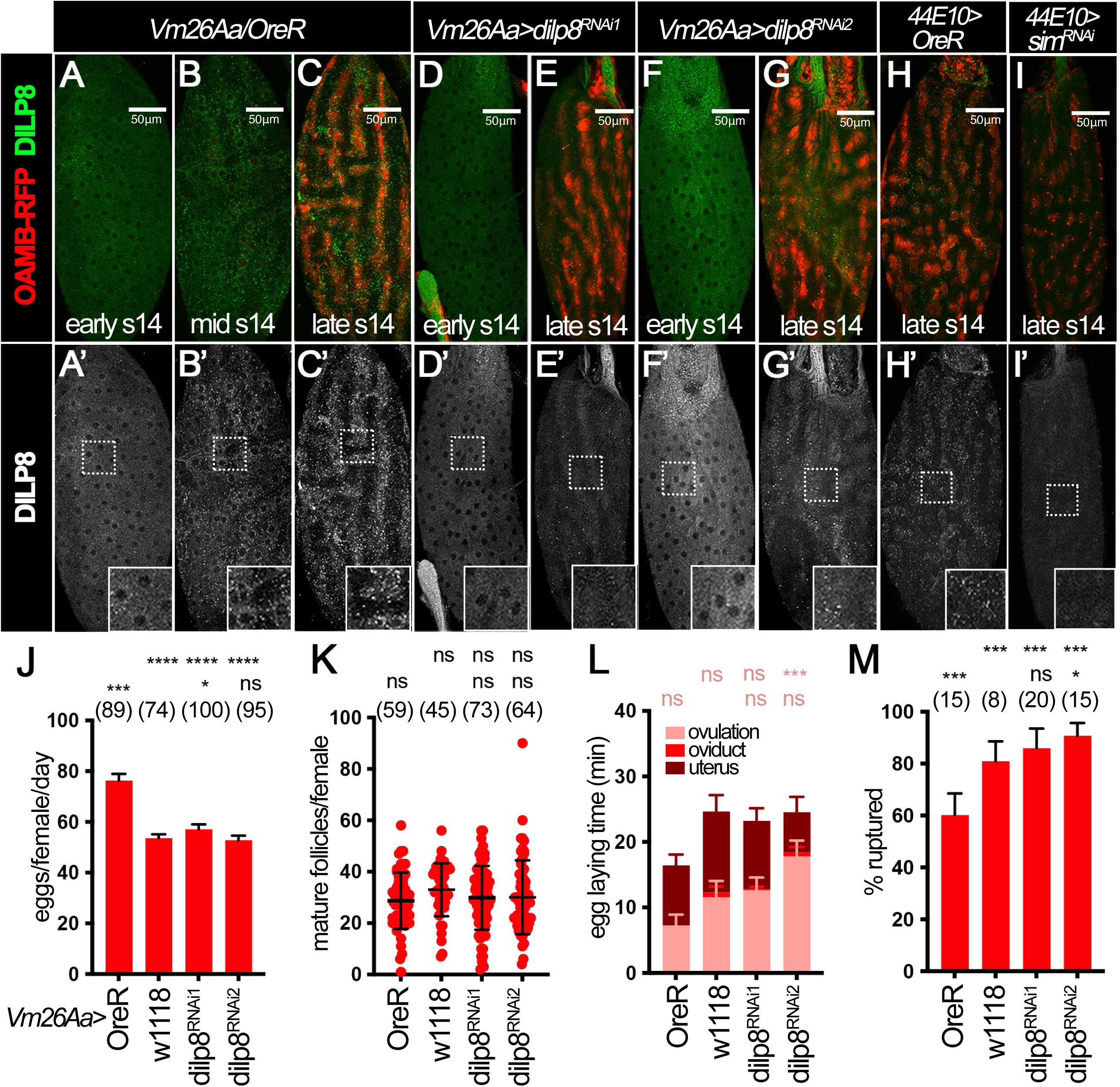
DILP8 is a downstream target of Sim and not required for mating-induced ovulation. **(A–C)** Expression of DILP8 (green in A-C and white in A’-C’) in early (A-A’), mid (B-B’), and late (C-C’) stage-14 follicles from control females. OAMB-RFP (red in A-C) is used to mark late stage-14 follicles. Insets show high magnification of squared area. **(D-G)** Expression of DILP8 (green in D-G and white in D’-G’) in stage-14 follicles from females with *Vm26Aa-Gal4* driving *dilp8^RNAi1^* (D-E’) or *dilp8^RNAi2^* (F-G’). **(H-I)** Expression of DILP8 (green in H-I and white in H’-I’) in late stage-14 follicles of control (H) and *sim^RNAi^* (I) females with *44E10-Gal4* and *Oamb-RFP*. **(J-L)** Quantification of egg laying (J), mature follicle numbers after egg laying (K), and egg laying time (L) in *Oregon R (OreR)* control, *w1118* control, *dilp8^RNAi1^, or dilp8^RNAi2^* females with *Vm26Aa-Gal4* and *Oamb-RFP*. The number of females is noted above each bar. **(M)** Quantification of OA-induced follicle rupture using mature follicles isolated from OreR, w1118, *dilp8^RNAi1^, or dilp8^RNAi2^* females with *Vm26Aa-Gal4* and *Oamb-RFP*. The number of wells analyzed (containing 25-35 follicles) is noted above each bar. For all bar graphs, data points represent mean ± SD, except egg laying graphs which depict mean + SEM. For egg laying time data, the significance depicted above each bar is for the ovulation time (also see Table S1). The statistical significance in the top row is relative to *OreR* control, and the bottom row is relative to *w1118* control. ***p<0.001, **p<0.01, *p<0.05, ns = not significant (One-way ANOVA). Scale bars are 50 μm.

Next, we examined DILP8 expression in *sim-knockdown* follicles using the DILP8 antibody. Consistent with the RNAseq analysis, DILP8 was not detected in mature follicles with *sim* knockdown (Figure 1H-I). Together, these data demonstrated that DILP8 is upregulated via Sim in mature follicle cells and suggested that DILP8 could play an important role in ovulation.

### DILP8 in mature follicle cells is not required for mating-induced ovulation and egg laying

Previous studies showed that *dilp8* homozygous mutants exhibit a significant reduction in egg laying capacity after mating and increased numbers of stage-10-14 egg chambers inside ovaries, which led authors to conclude that DILP8 regulates ovulation and female fecundity (Li et al., 2023; Liao and Nässel, 2020). To confirm these studies, we examined whether *dilp8* knockdown using *Vm26Aa-Gal4* driver in follicle cells also leads to an ovulation defect. We included two experimental controls, one with wild-type *Oregon R* (*w+*) and the other with a *white* mutation (*w^1118^*), the latter of which has the same genetic background as the RNAi lines. To our surprise, *dilp8*-knockdown females laid more than 50 eggs/female/day after mating with wild-type males, which had no significant difference from control females with *w^1118^* background (Figure 1J). We did notice that the control with a wild-type *Oregon R* background showed significantly higher egg laying capacity, which could be due to different genetic background. Since the control with *w^1118^* is more appropriate for RNAi lines, as they have the same genetic background, we concluded that *dilp8* knockdown in follicle cells did not lead to defects in mating-induced egg laying. In addition, we also measured the number of mature follicles retained in the ovaries after egg laying and calculated the time required for ovulation, two major indicators of an ovulation defect (Beard et al., 2023). Consistent with the egg-laying assay, *dilp8*-knockdown females showed a normal number of mature follicles inside the ovaries after egg laying (Figure 1K) and spent a normal amount of time ovulating eggs (Figure 1L and Table S1). Furthermore, we performed the *ex vivo* follicle rupture assay, which we previously developed to directly evaluate the ovulatory competency of the mature follicles in response to the ovulatory stimulus octopamine (OA; Beard et al., 2023; Deady and Sun, 2015; Knapp et al., 2018). Consistent with all other experiments, *dilp8*-knockdown mature follicles could undergo normal (or even better) follicle rupture in response to OA (Figure 1M and S1E-H), indicating that *dilp8*-knockdown follicles are fully matured and ready for ovulation. Altogether, our data suggest that DILP8 in mature follicles is not required for mating-induced ovulation and egg laying.

### Global expression of DILP8 is not required for mating-induced ovulation and egg laying

The difference between our experiments and previous reports could be that we only knock down *dilp8* in follicle cells or due to an incomplete *dilp8* knockdown. To test this hypothesis, we utilized the following three *dilp8* mutant lines to deplete *dilp8* globally (Figure 2A): *dilp8^MI00727^* (*dilp8^MI^*), a strong loss-of-function allele with a MiMIC insertion with GFP coding sequence to disrupt *dilp8* splicing (Liao and Nässel, 2020; Venken et al., 2011); *dilp8^EX^*, a small deletion that partially truncates the 5’ *dilp8* gene and neighboring genes (Colombani et al., 2012); and *Df(3L)BSC414* (*BSC414*), a large depletion that affects 62 genes including *dilp8* (information from FlyBase). Using the DILP8 antibody, we first validated that DILP8 is not detected in mature follicles from *dilp8^EX/MI^*, *dilp8^EX/EX^*, and *dilp8^EX^/BSC414* females (Figure 2B-E). Next, we performed the mating-induced egg laying, mature follicle retention after egg laying, egg laying time, and OA-induced follicle rupture using these *dilp8* mutant females. In comparison to the *dilp8^EX^/w^1118^* heterozygous background, all *dilp8* mutant females showed no defects in mating-induced egg laying (Figure 2F and Table S1), ovulation time (Figure 2H and Table S1), and OA-induced follicle rupture (Figure 2I, S1I-L). We did observe a slight but statistically significant increase in mature follicles in *dilp8^EX/MI^* and *dilp8^EX^/BSC414* females and a decrease in mature follicles in *dilp8^EX/EX^* females (Figure 2G). Altogether, these data suggest that global *dilp8* depletion does not result in an ovulation defect in the mated condition, which is inconsistent with previous reports.

**Figure 2.**
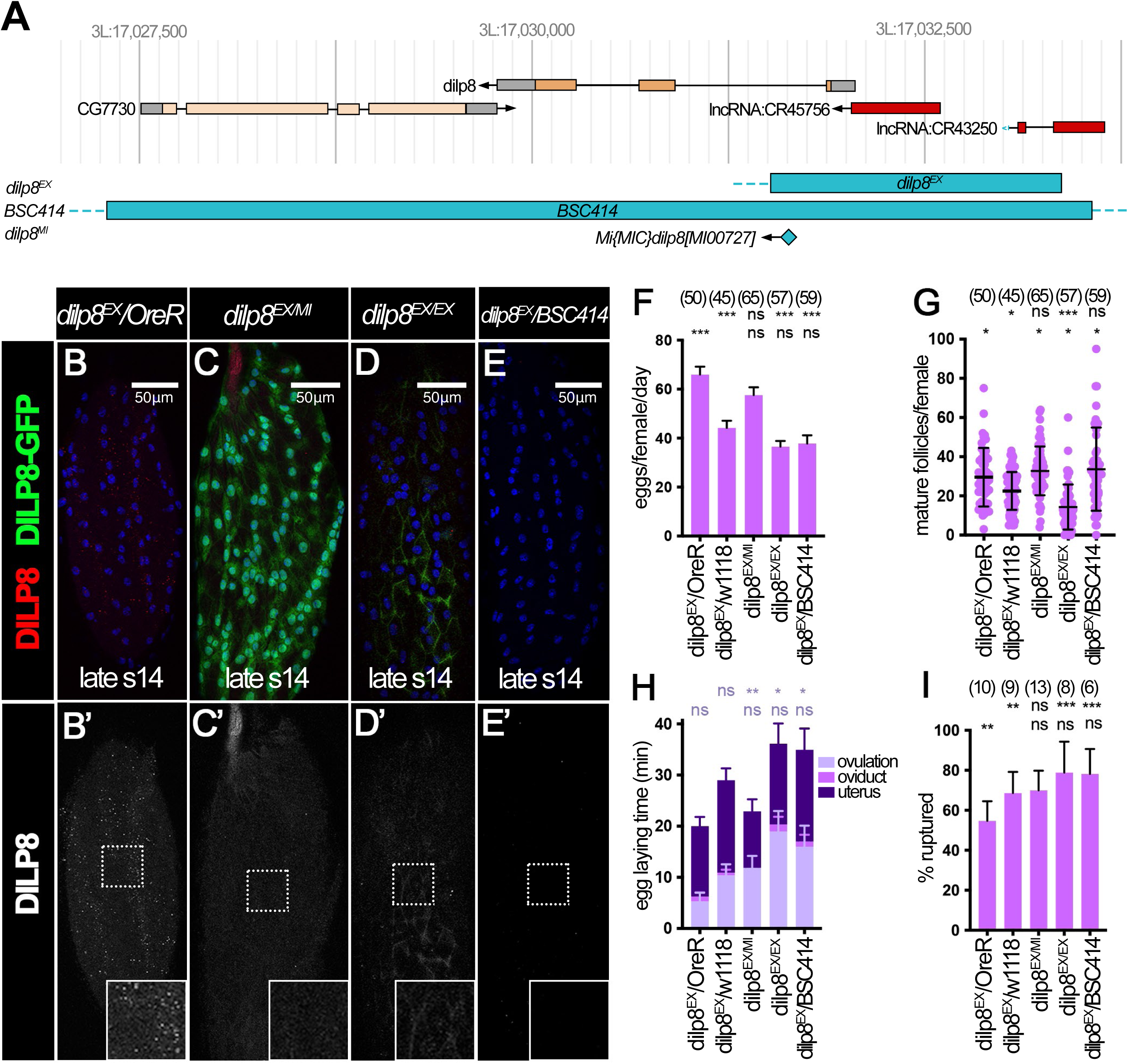
*dilp8* mutant females have no defects in mating-induced ovulation and follicle rupture. **(A)** A schematic depicts the nature of *dilp8* mutant alleles. **(B-E)** Expression of DILP8 (red in B-E and white in B’-E’) in late stage-14 follicles from *dilp8^EX^/Oregon-R* (B-B’), *dilp8^EX/MI^* (C-C’), *dilp8^EX/EX^* (D-D’), and *dilp8^EX^/BSC414* (E-E’) females. *dilp8^MI^* contains GFP reporter (green in C) reflecting *dilp8* transcription activity (Liao and Nässel, 2020). Insets show higher magnification of squared area. All images from B-E were acquired using the same microscopic settings. Nuclei are marked by DAPI in blue. Scale bar is 50um. **(F-H)** Quantification of egg laying (F), mature follicle numbers after egg laying (G), and egg laying time (H) in *dilp8^EX^/OreR* heterozygous control, *dilp8^EX^/w1118* heterozygous control, *dilp8^EX/MI^*, *dilp8^EX/EX^* or *dilp8^EX^/BSC414* mutant females. The number of females is noted above each bar. **(I)** Quantification of OA-induced follicle rupture using mature follicles isolated from *dilp8^EX^/OreR* heterozygous control, *dilp8^EX^/w1118* heterozygous control, *dilp8^EX/MI^*, *dilp8^EX/EX^* or *dilp8^EX^/BSC414* mutant females. The number of wells analyzed (containing 25-35 follicles) is noted above each bar. For all bar graphs, data points represent mean ± SD, except egg laying graphs which depict mean + SEM. For egg laying time data, the significance depicted above each bar is for the ovulation time (also see Table S1). The statistical significance in the top row is relative to *OreR* control, and the bottom row is relative to *w1118* control. ***p<0.001, **p<0.01, *p<0.05, ns = not significant (One-way ANOVA).

### Ovarian DILP8 is required to prevent excessive accumulation of mature follicles in virgin females

The lack of an ovulation and egg laying defect in *dilp8* mutant females left us wondering what the function of DILP8 in mature follicles was. In an exploratory experiment, we dissected the ovaries from control and *dilp8* mutant females fed with wet yeast for 9 days and discovered that the ovarioles of the *dilp8* mutant females contain more than three (seven in an extreme case) mature follicles, which is seldom observed in control females (Figure S2A-C). This preliminary data prompted us to hypothesize that DILP8 functions as a mature follicle sensor to prevent excessive accumulation of mature follicles inside the ovaries. To test this hypothesis, we utilized the virgin females to examine the follicle composition in each ovariole (Figure 3A) because virgin females tend to accumulate mature follicles inside their ovaries without mating-induced ovulation. We first analyzed the ovariole distribution from control virgin females fed with wet yeast for three or nine days. On average, each control ovariole has 4.67 ± 0.57 stage-1-7 follicles, 0.79 ± 0.41 stage-8-9 follicles, 0.25 ± 0.43 stage-10 follicles, 0.18 ± 0.38 stage-11-13 follicles, and 1.67 ± 0.57 stage-14 follicles in the 3-day-feeding condition (Figure 3B and E, S3A). Similarly, each control ovariole in the 9-day-feeding condition has 4.93 ± 0.48 stage-1-7 follicles, 0.72 ± 0.47 stage-8-9 follicles, 0.20 ± 0.40 stage-10 follicles, 0.10 ± 0.31 stage-11-13 follicles, and 1.38 ± 0.55 stage-14 follicles (Figure 3F, S3B). In contrast, the ovarioles from *dilp8* mutant females have significantly more stage-14 follicles in both 3-day-feeding and 9-day-feeding conditions, except *dilp8^EX/EX^* mutant in the 3-day-feeding condition (Figure 3C-F and S3A-B). In addition, there is an increased percentage of ovarioles with more than three stage-14 follicles (Figure 3E-F). These data suggest that DILP8 is required to prevent excessive accumulation of mature follicles in virgin females.

**Figure 3.**
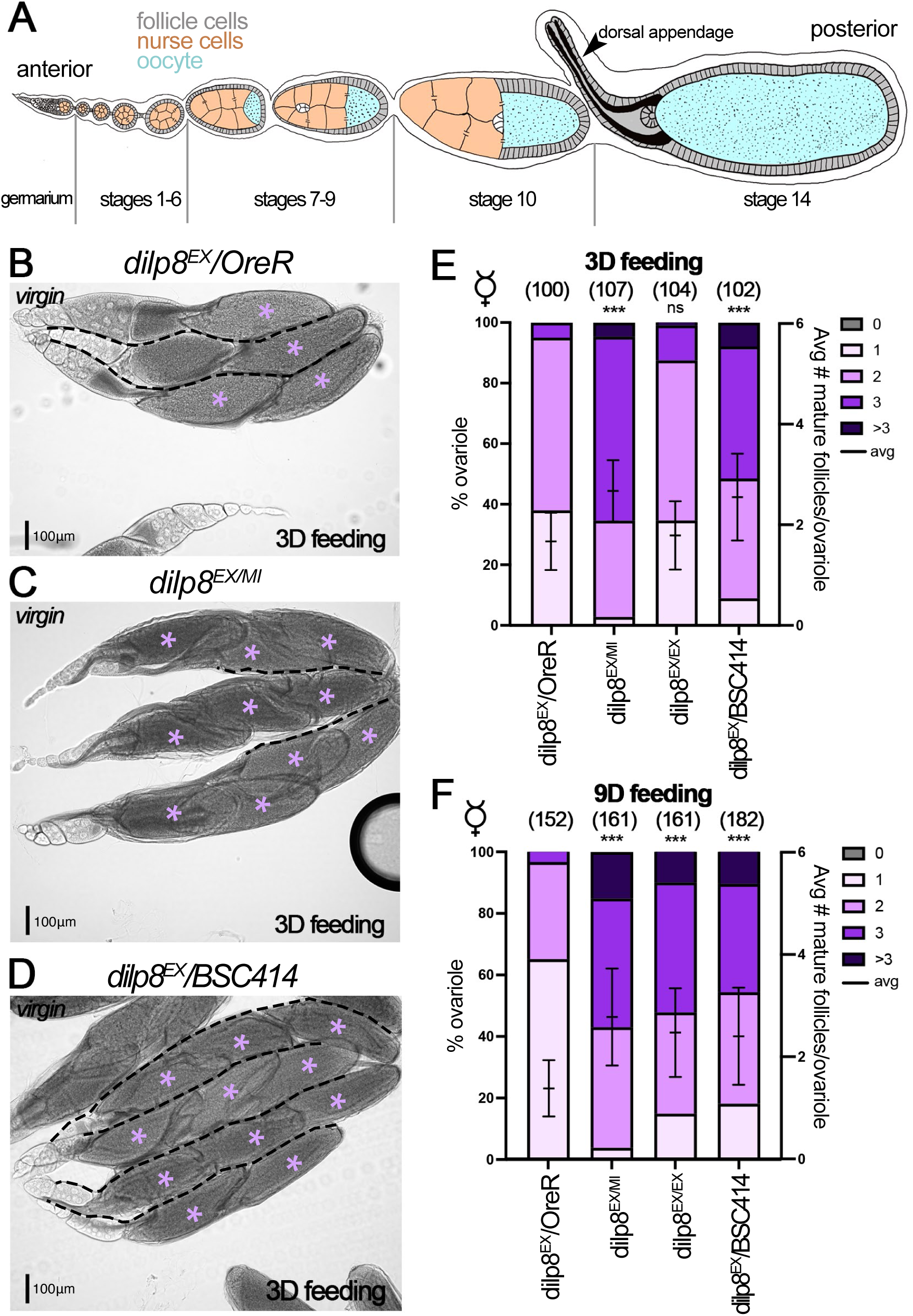
Global depletion of *dilp8* results in an increased number of mature follicles in virgin females. **(A)** An illustration depicts the ovariole with the germarium at the anterior and subsequent stages leading to stage-14 follicle at the posterior. The dorsal appendage structure is labeled with an arrowhead. **(B-D)** Representative brightfield images of ovarioles from *dilp8^EX^/OreR* heterozygous control (B), *dilp8^EX/MI^* (C) or *dilp8^EX^/BSC414* (D) virgin females after three-day wet-yeast feeding. Mature follicles are denoted with purple asterisks and ovarioles are separated by black dashed lines when needed. The scale bar is 100 μm. **(E-F)** Quantification of mature (stage-14) follicle per ovariole in *dilp8^EX^/OreR* heterozygous control, *dilp8^EX^/w1118* heterozygous control, *dilp8^EX/MI^*, *dilp8^EX/EX^* or *dilp8^EX^/BSC414* virgin mutant females after three-day (E) or nine-day (F) wet-yeast feeding. The number of ovarioles is noted above each bar. The left y-axis represents the percentage of ovarioles with 0, 1, 2, 3, or >3 mature follicles, and the right y-axis represents the average number of mature follicles per ovariole, denoted by a line with standard deviation. Significance relative to the *OreR* controls is provided above each bar. ***p<0.001, **p<0.01, *p<0.05, ns = not significant (One-way ANOVA).

To determine whether the ovarian DILP8 is responsible for preventing excessive accumulation of stage-14 follicles, we also evaluated the ovariole distribution in *dilp8*-knockdown virgin females with *Vm26Aa-Gal4* driver. Consistent with *dilp8* mutant females, ovarioles of *dilp8*-knockdown virgin females have significantly more stage-14 follicles in both 3-day- and 9-day-feeding conditions (Figure 4A-D and S3C-D). Altogether, our data indicated that ovarian DILP8 is required to prevent the excessive accumulation of mature follicles in virgin females.

**Figure 4.**
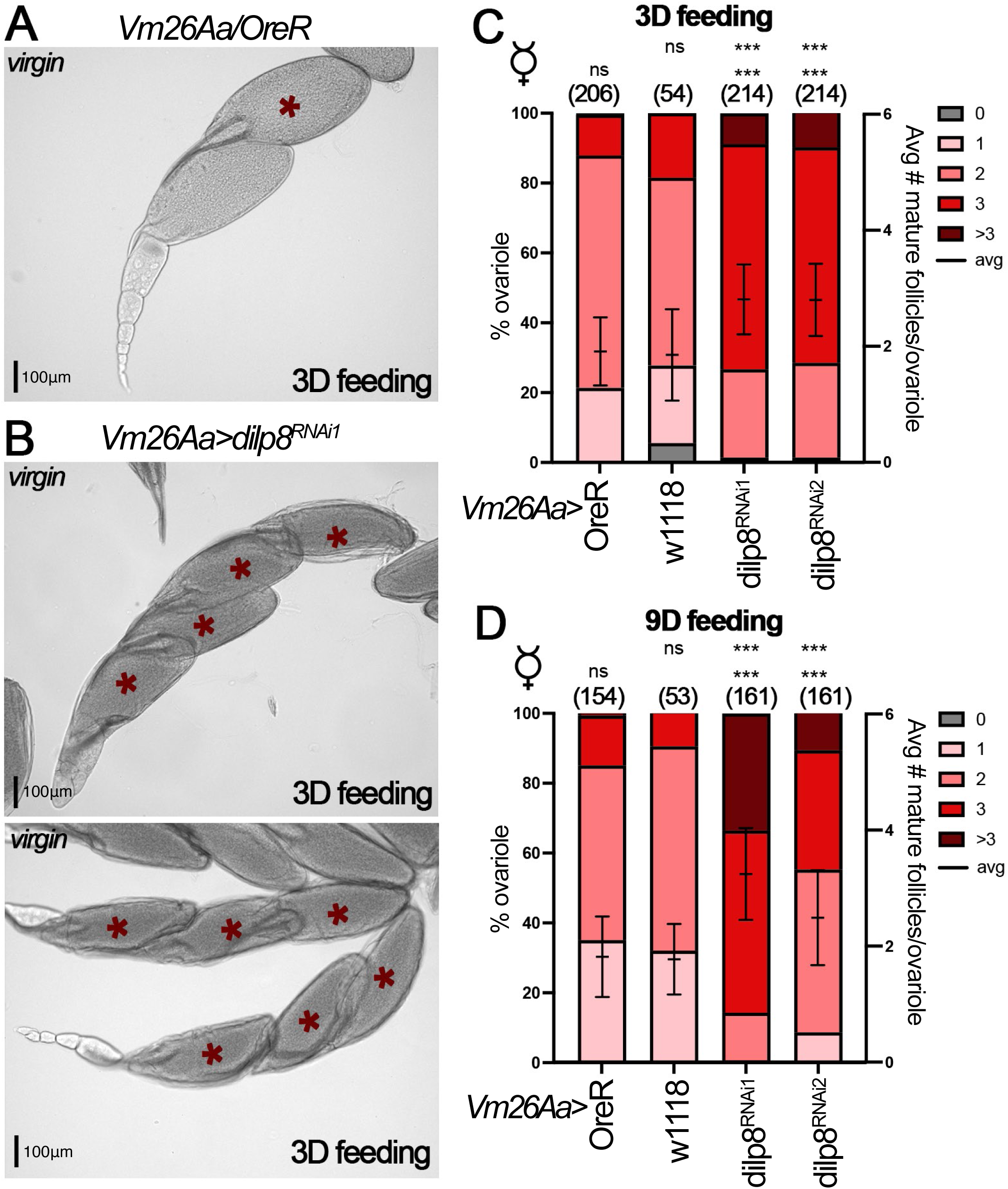
Ovarian depletion of *dilp8* results in an increased number of mature follicles in virgin females. **(A-B)** Representative brightfield images of ovarioles from *OreR* control (A) or *dilp8^RNAi1^*(B) virgin females with *Vm26Aa-Gal4* after three-day wet-yeast feeding. Mature follicles are denoted with red asterisks. The scale bar is 100 μm. **(C-D)** Quantification of mature (stage-14) follicle per ovariole in *OreR* control, *w1118* control*, dilp8^RNAi1^*, or *dilp8^RNAi2^* virgin females (with *Vm26Aa-Gal4* and *Oamb-RFP*) after three-day (C) or nine-day (D) wet-yeast feeding. The number of ovarioles is noted above each bar. The left y-axis represents the percentage of ovarioles with 0, 1, 2, 3, or >3 mature follicles and the right y-axis represents the average number of mature follicles per ovariole, denoted by a line with standard deviation. The statistical significance in the top row is relative to *OreR* controls, and the bottom row is relative to *w1118* controls. ***p<0.001, **p<0.01, *p<0.05, ns = not significant (One-way ANOVA).

### Ovarian DILP8 is required for ovulation in virgin females to prevent oocyte aging

The increase in the number of mature follicles per ovariole in *dilp8*-depleted virgin females could be due to defects in virgin ovulation. To test this hypothesis, we first evaluated whether *dilp8* mutant virgin females have an egg-laying defect. We performed a modified egg-laying assay using virgin *dilp8* mutant females (see materials and methods). Consistent with our hypothesis, all three *dilp8* mutant virgin females (*dilp8^EX/MI^*, *dilp8^EX/EX^*, *dilp8^EX^/BSC414*) laid less than half of the eggs in comparison to control virgin females over 10 days (Figure 5A-A’). In addition, females with *dilp8* knockdown in follicle cells showed similar defects in virgin egg laying (Figure 5B-B’). To evaluate if the virgin egg laying defect is caused by an ovulation defect or another step in the egg-laying process (such as egg transport through the oviduct and oviposition; Beard et al., 2023), we quantified the frequency of eggs located in each part of the reproductive tract in females dissected after completion of the 10-day virgin egg laying assay. We observed that after egg laying, ovarian *dilp8*-knockdown virgin females did not show a higher frequency of follicles in the oviduct or uterus (Figure S4), suggesting that the reduction in virgin egg laying is unlikely due to defects in egg transport or oviposition. Altogether, our data indicate that ovarian DILP8 is required for ovulation in virgin females to prevent excessive accumulation of mature follicles.

**Figure 5.**
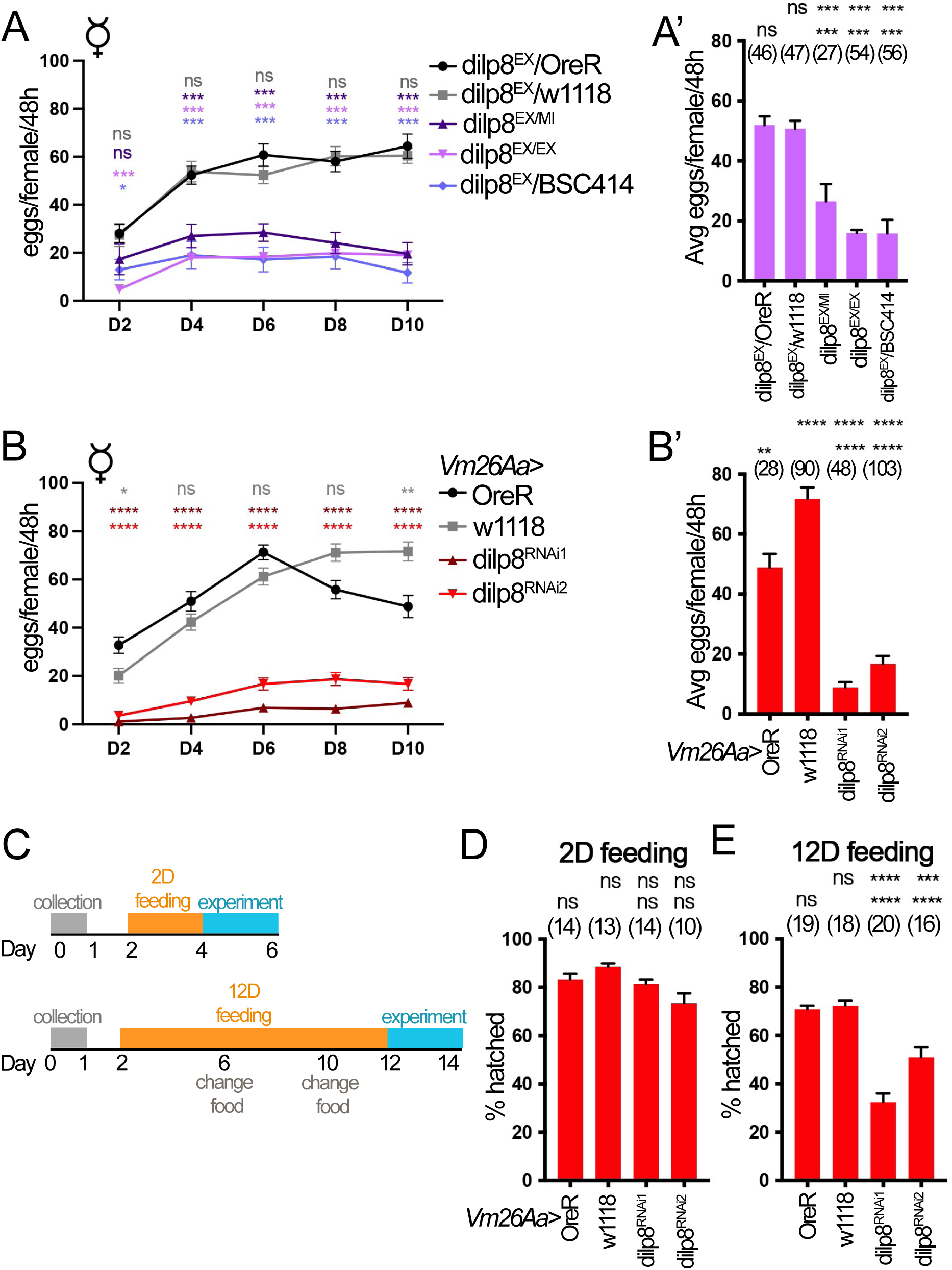
Ovarian DILP8 is required for ovulation in virgin females to prevent oocyte aging. (A-B’) Quantification of laid eggs per virgin female every 48 hours over ten days. Line graphs (A and B) show the average number of eggs laid per female every 48 hours at each specific day for multiple genotypes with the significance value denoted at the top of each data point, color-coded for each genotype. The corresponding bar graphs (A’and B’) show the average number of eggs laid per female per 48h over the 10 days with the total number of females depicted above each bar along with significance. Datasets for *dilp8^EX^/OreR* heterozygous control, *dilp8^EX^/w1118* heterozygous control*, dilp8^EX/MI^*, *dilp8^EX/EX^* or *dilp8^EX^/BSC414* virgins (A-A’). Datasets for *OreR* control, *w1118* control*, dilp8^RNAi1^*, or *dilp8^RNAi2^* virgin females with *Vm26Aa-Gal4* and *Oamb-RFP* (B-B’). **(C)** Schematic of experimental timeline and description of feeding during hatchability experiments. **(D-E)** Quantification of the percentage of eggs hatched for 2-day-fed (D) or 12-day-fed (E) virgin females after mating with fertile *OreR* males for 24 hours. The number of females is depicted above each bar. Significance relative to *OreR* controls is provided above each bar on the top row, whereas significance relative to the *w1118* is on the bottom row. For all bar graphs, data points represent mean ± SD, except egg laying graphs which depict mean ± SEM. ***p<0.001, **p<0.01, *p<0.05, ns = not significant (One-way ANOVA).

Previous work showed that artificially blocking egg laying leads to decreased oocyte quality and oocyte aging, manifested by reduced hatching rates (hatchability) of the laid eggs (Greenblatt et al., 2019). Therefore, we reasoned that excessive accumulation of mature follicles in *dilp8*-knockdown virgin females could lead to reduced oocyte quality and oocyte aging. To test this hypothesis, we fed virgin females for either 2 days (no mature follicle retention) or 12 days (with mature follicle retention) and then mated with fertile males to evaluate the quality of the retained oocytes (Figure 5C). Consistent with our expectation, eggs from 2-day-fed *dilp8*-knockdown females had normal hatchability (Figure 5D), while eggs from 12-day-fed *dilp8*-knockdown females had significantly lower hatchability than controls (Figure 5E). Therefore, ovarian DILP8 is required for ovulation in virgin females to prevent the excessive accumulation of mature follicles and oocyte aging.

### Lgr3 is likely a receptor to mediate ovarian DILP8’s effect

Since Lgr3 is the receptor for DILP8 in regulating organ size and developmental timing (Colombani et al., 2015; Garelli et al., 2015; Vallejo et al., 2015), we tested whether Lgr3 is the receptor to mediate ovarian DILP8’s effect on inducing virgin ovulation and preventing excessive accumulation of mature follicles. Females with global *Lgr3* knockdown with *act-Gal4* driver showed normal mating-induced egg laying and ovulation time (Figure 6A and C). The mature follicle counts after egg laying are normal in *Lgr3^RNAi1^* but were slightly higher in *Lgr3^RNAi2^*. In contrast, *Lgr3*-knockdown virgin females showed a significant increase of mature follicles in both 3-day and 9-day feeding conditions (Figure 6D-E). In addition, *Lgr3*-knockdown virgin females laid significantly fewer eggs than control virgin females (Figure 6F-F’). Notably, *Lgr3^RNAi2^*showed more severe defects than *Lgr3^RNAi1^*, consistent with previous reports that *Lgr3^RNAi2^* (using the V22 vector) is more efficient to knock down *Lgr3* in development (Garelli et al., 2015). As expected, eggs from 2-day-fed *lgr3*-knockdown females had normal hatchability (Figure 6G), while eggs from 12-day-fed *Lgr3^RNAi2^* females had significantly lower hatchability than controls (Figure 6H). Altogether, loss of *Lgr3* mimics the *dilp8* mutant phenotype, suggesting that Lgr3 is likely the receptor for ovarian DILP8 in regulating virgin ovulation. In conclusion, ovarian DILP8, via Lgr3, functions as a mature follicle sensor to prevent excessive accumulation of mature follicles and oocyte aging in virgin females by promoting virgin ovulation (Figure 7).

**Figure 6.**
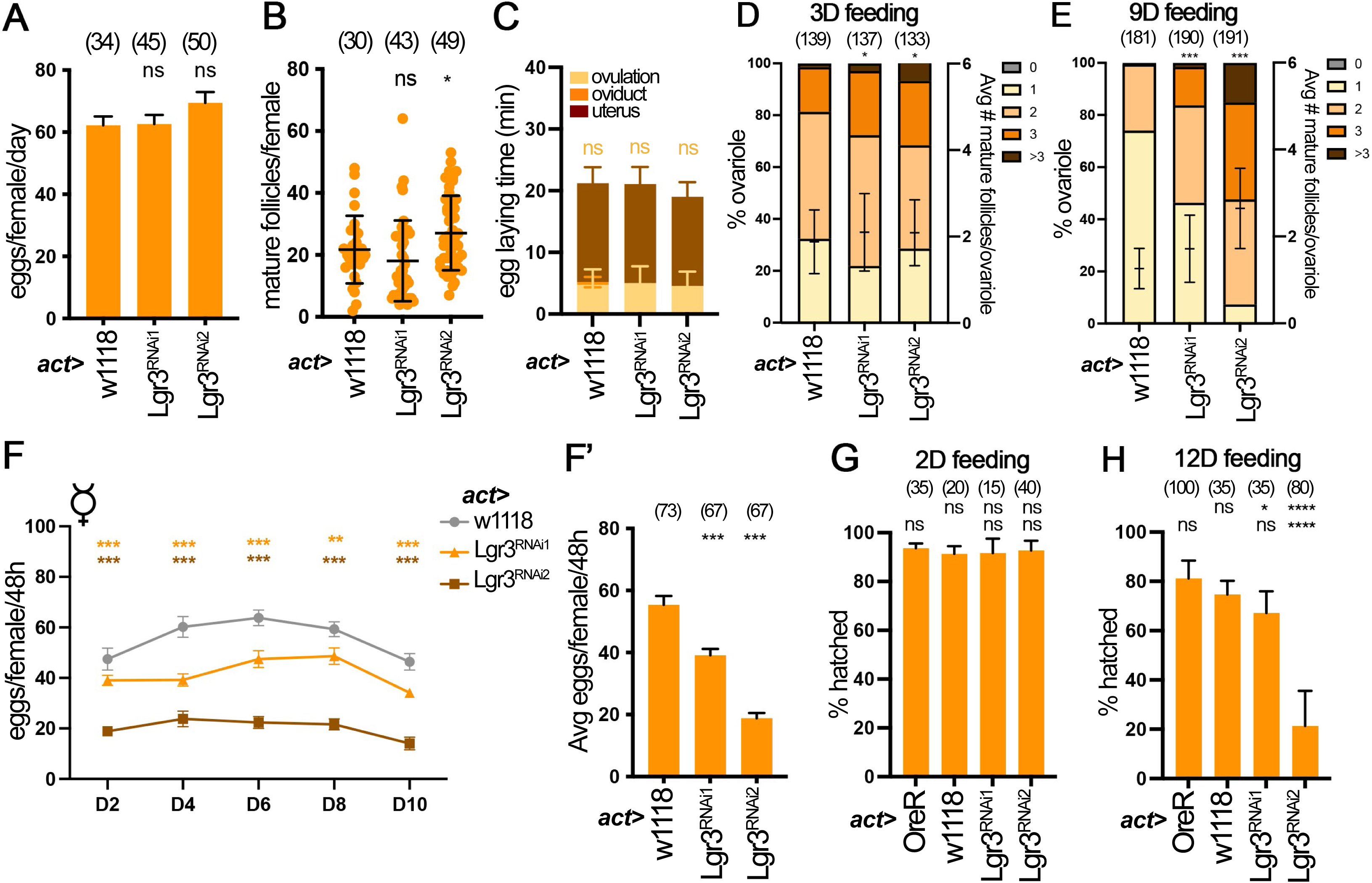
Lgr3 is likely a receptor to mediate ovarian DILP8’s effect. (A-C) Quantification of egg laying (A), mature follicle numbers after egg laying (B) and egg laying time (C) in *w1118* control, *Lgr3^RNAi1^*, or *Lgr3^RNAi2^* mated females with *act-Gal4, tub-gal80ts*. The number of females is noted above each bar. **(D-E)** Quantification of mature (stage-14) follicle numbers per ovariole after three-day (D) or nine-day (E) wet-yeast feeding for *w1118* control, *Lgr3^RNAi1^*, or *Lgr3^RNAi2^* virgin females with *act-Gal4, tub-gal80ts*. The number of ovarioles is noted above each bar. The left y-axis represents the percentage of ovarioles with 0, 1, 2, 3, or >3 mature follicles and the right y-axis represents the average number of mature follicles per ovariole, denoted by a line with standard deviation error bars. **(F)** Quantification of laid eggs per virgin female every 48h over 10 days. The line graph (F) shows the average number of eggs laid per female every 48h on a specific day. The significance value is denoted at the top of each data point color-coded for each genotype. A corresponding bar graph (F’) shows the average number of eggs laid per female per 48h over the 10 days. **(G-H)** Quantification of the percentage of eggs hatched for 2-day-fed (G) or 12-day-fed (H) virgin females after mating with fertile males for 24 hours. The number of females is depicted above each bar. Significance relative to *OreR* controls is provided above each bar on the top row, whereas significance relative to the *w1118* is on the bottom row. For all bar graphs, data points represent mean ± SD, except egg laying graphs which depict mean ± SEM. ***p<0.001, **p<0.01, *p<0.05 (One-way ANOVA).

**Figure 7.**
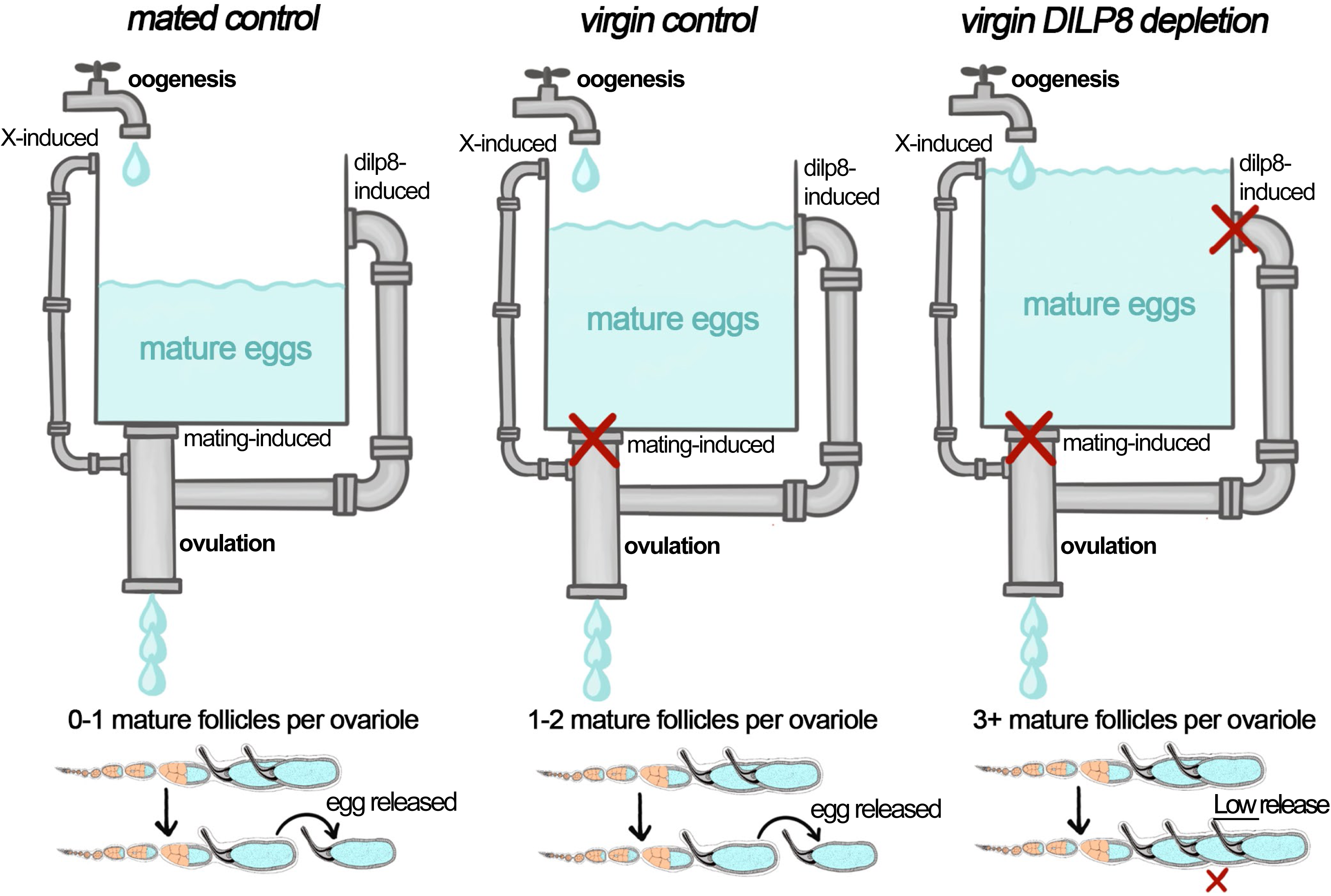
DILP8 acts as a mature follicle sensor to prevent excessive accumulation of mature follicles in *Drosophila* ovaries. Schematic model of the function of DILP8 in mated and virgin female ovulation in control versus DILP8-depleted females. This sink model metaphorically represents the process. In the model, water represents the number of mature follicles, and oogenesis (egg production) is depicted by an open faucet (assuming a constant rate for simplification). The three pipes represent the ovulation processes that can take place: mating-induced, dilp8-induced (mating-independent), and ovulation induced by an unknown X-factor (mating-independent). The width of each pipe is meant to depict that some ovulation processes may result in a larger number of mature eggs released relative to others.

## Discussion

In this work, we found that the number of mature follicles in wild-type *Drosophila* ovaries is tightly controlled, with 0-2 mature follicles per ovariole regardless of the female’s mating status (Figures 3 and 4). In the mated condition, mating induces an increase in ovulation rate as well as the number of germline stem cells (Yoshinari et al., 2020), the latter of which presumably leads to an increase in follicle production. This likely balances the oogenesis and ovulation rate to ensure the constant number of mature follicles in the ovaries (Figure 7). However, in virgin conditions, the mechanism by which the ovary counts the number of mature follicles to coordinate oogenesis and ovulation has been a mystery in the reproductive field, and it is not fully understood how the ovariole can constantly maintain this precise number of mature follicles. In addition, it is unknown how virgin females prevent excessive long-term storage of their mature follicles to ensure the high quality of their stored oocytes before the next mating. This work contributes to this knowledge gap by uncovering DILP8 as the mating-independent signal that allows the females to sense the number of mature follicles in their ovaries and get rid of excessive mature follicles by inducing ovulation. The increase of mature follicles in virgin females will increase DILP8 levels, which likely activates the Lgr3 receptor to induce ovulation and release the oldest mature follicle in each ovariole when reaching the threshold level (Figure 7). In the absence of DILP8 or the Lgr3 receptor, the number of mature follicles will keep increasing and the quality of their oocytes will decrease, which also provides a novel assay to study oocyte aging in the future (Greenblatt et al., 2019). Therefore, DILP8 functions as a mature follicle sensor to ensure the homeostasis of mature follicles in the female ovaries. This is very similar to DILP8’s role in development as an organ size sensor to ensure proper development of adult organ size. It is intriguing to generalize that DILP8/Lgr3 functions as a homeostatic regulator of multiple developmental and physiological processes in *Drosophila* and beyond, which awaits further confirmation by future discoveries.

To this date, there was no information regarding how virgin ovulation was induced. Our discovery that DILP8 and Lgr3 play an essential role in virgin ovulation can act as a foundation to expand our knowledge about this mechanism in the future. Unlike previous reports (Liao and Nässel, 2020), we cannot detect any Lgr3 expression in the ovaries or posterior mature follicles using the Lgr3::GFP reporter. Therefore, we speculate that DILP8 must be secreted into the hemolymph to reach the Lgr3 receptor in extra ovarian tissues, most likely the sexually dimorphic Lgr3+ neurons (Liao and Nässel, 2020; Meissner et al., 2016), which will be in our future investigation. It is also worth noting that DILP8/Lgr3 is unlikely the only signal to regulate virgin ovulation as *dilp8* mutation did not completely block virgin ovulation and egg laying (Figure 5A). Therefore, there is another backup mechanism to induce virgin ovulation in addition to DILP8, and the nature of this mechanism is unknown (Figure 7). Since octopaminergic neurons are critical for mating-induced ovulation and octopamine works directly on mature follicle cells for follicle rupture and ovulation (Deady and Sun, 2015; White et al., 2021), it will also be interesting to test whether octopaminergic neurons play a role in virgin ovulation.

In addition, our work also revealed that transcription factor Sim acts upstream to regulate DILP8 expression in mature follicle cells. It is unknown whether DILP8 is a direct target of Sim. Contrary to our prediction and previous findings, we found that DILP8 is not essential for regulating the ovulatory competency of mature follicles and for mating-induced ovulation and egg laying in optimal conditions. Previous works (Li et al., 2023; Liao and Nässel, 2020) showed that *dilp8^MI/MI^* homozygous mutant females laid significantly fewer eggs and accumulated more mature follicles in their ovaries than control females after mating, which led authors to conclude that DILP8 is required for mating-induced ovulation. First, it is unknown whether *dilp8^MI^* stocks accumulate background mutations that contribute to the observed phenotype, as we were unable to find *dilp8^MI/MI^* homozygous flies in our stocks. Second, the average number of eggs laid by control females in their studies is around 20/female/day, which is way lower than what we observed in our optimal experimental conditions (about 60/female/day). This leads to the speculation that the experimental condition in the previous studies may be not optimal for egg laying, which leads to the accumulation of mature follicles in both control and *dilp8^MI/MI^* females, mimicking the virgin condition in our experimental setting. Unfortunately, we cannot repeat the previous experiments since we were unable to get *dilp8^MI/MI^* homozygous females.

The role of ILP8/Lgr3 in regulating ovulation is not limited to *Drosophila*. Homologs of ILP8 (Gonadulin) and Lgr3 are found in the kissing bug *Rhodnius prolixus* and play critical roles in ovulation after blood meal feeding and mating (Leyria et al., 2022). It is unclear whether *Rhodnius* Gonadulin/Lgr3 is required for virgin ovulation as in *Drosophila*. In addition, Gonadulin is also found to be specific in locust ovaries, but its role in ovulation has not been determined (Veenstra et al., 2021). According to a recent study, tumor-derived DILP8/INSL3 plays conserved roles in inducing cancer anorexia via Lgr3/Lgr8 in both *Drosophila* and mammals (Yeom et al., 2021). INSL3 (Insulin-like peptide 3) is also produced in mammalian ovaries, particularly theca cells of growing antral follicles, and likely regulates follicle maturation and ovulation (Ivell and Anand-Ivell, 2018). Our findings in *Drosophila* lead to a prediction that mammalian INSL3 may be playing a similar role in sensing the number of antral follicles to coordinate primordial follicle recruitment and ovulation, which would be advantageous to prevent unnecessary depletion of the ovarian reserve and avoidable energy consumption for excess follicle maintenance. Future work is required to test this hypothesis.

## Materials & Methods

### Drosophila genetics

Except otherwise noted, *Drosophila melanogaster* flies were reared on standard cornmeal-molasses food at 25 °C. All RNAi-mediated depletion experiments had *UAS-dcr2* and were shifted to 29 °C upon adult fly eclosion to increase knockdown efficiency. The following Gal4 drivers were used: *44E10-Gal4* from the Janelia Gal4 collection (Deady and Sun, 2015; Pfeiffer et al., 2008), *Vm26Aa-Gal4* (Peters et al., 2013), and *act-Gal4/Cyo; tub-Gal80ts/TM3*. To knockdown genes with above mentioned Gal4 drivers, the following RNAi lines were used: *UAS-dilp8^RNAi1^* (Vienna *Drosophila* Resource Center - VDRC, stock #102604), *UAS-dilp8^RNAi2^* (VDRC, stock #9420), *UAS-sim^RNAi^* (VDRC, stock #26888), UAS-Lgr3^RNAi1^ (Bloomington *Drosophila* Stock Center - BDSC, stock #55910), Lgr3^RNAi2^ (BDSC, stock #36887). The following mutant lines were used for dilp8 mutant analysis: *dilp8^MI00727^/TM3, Sb, Ser* (BDSC, stock #33079), *Df(3L)BSC414/TM6C, Sb, cu* (BDSC, stock #24918), *dilp8^EX^/TM3,Sb* (Colombani et al., 2012). Isolation and identification of stage-14 follicles for follicle rupture assay were performed using *Oamb-RFP* (Knapp et al., 2019). Control flies for all experiments were prepared by crossing Gal4 driver to *Oregon-R (w+)* or *w^1118^* flies. For the generation of flip-out Gal4 clones, the *hsFLP;; act<CD2<Gal4, UAS-GFP/TM3* stock was crossed to the indicated transgenes of interest. Adult female progeny with correct genotypes were heat shocked for 45 min at 37°C for clone induction and subsequently incubated at 25 °C for 2-4 days before dissection.

### Immunostaining and microscopy

The immunostaining procedure was carried out similarly to what has been previously reported (Oramas et al., 2023) with minor modifications. In short, ovaries were dissected in cold Grace’s medium and fixed in 4% EM-grade paraformaldehyde in PBT (1XPBS + 0.2% Triton X-100) for 20 minutes at room temperature. The tissue was washed vigorously with PBT, blocked in antibody buffer (PBT + 0.5% BSA+ 2% normal goat serum), and stained with primary antibodies overnight at 4°C, followed by secondary antibody staining for 2 hours at room temperature. Ovaries were incubated with DAPI (final concentration 0.5 ug/mL) for 15 minutes before mounting. The following primary antibodies were used: rabbit anti-GFP (1:4000; Invitrogen, #A11122), rabbit anti-RFP (1:2000, MBL International, #PM005); rabbit anti-DILP8-A2 (1:2500 ; targeting DILP870−84; Liao and Nässel, 2020). Alexa Fluor 488 and Alexa Fluor 568 goat secondary antibodies (1:1000; Invitrogen, #A11001, #A11008, #A11004, #A11011) were used. Fluorescent and bright field polarized images were acquired using a Leica TCS SP8 confocal microscope and assembled using Adobe Photoshop software and ImageJ/Fiji software (https://fiji.sc/).

### Ovulation, hatchability, and follicle rupture assays

The mated egg laying, mature follicle count, and egg laying time assays were conducted and analyzed as described in previous literature (Beard et al., 2023; Deady and Sun, 2015; Deady et al., 2015; Knapp and Sun, 2017). In short, five- to six-day-old virgin females fed with wet yeast for a day were used to evaluate the number of eggs laid over two days after mating with *Oregon-R* males, or to examine the location of eggs in the reproductive tract after six-hour mating with *Oregon-R* males. These data were then used to calculate the average number of eggs laid/female/day and the average time required to ovulate an egg. Ovaries of females were dissected and evaluated after egg laying to count the number of mature follicles present in each female.

The *ex vivo* OA-induced follicle rupture assay was also conducted as previously described (Beard et al., 2023; Deady and Sun, 2015; Knapp et al., 2018). In this assay, five- to six-day-old virgin females are fed with wet yeast for three days. Their ovaries are dissected out and 25-35 intact stage-14 follicles are isolated using the mature follicle OAMB-RFP reporter, and incubated in a culture medium with OA (20 µM, Sigma-Aldrich) for 3 hours at 29 °C. The percentage of follicles ruptured after culture is quantified.

For the virgin egg laying assay, one- to two-day-old virgin females were fed with wet yeast for 2-days before the experiment. Five virgin females were placed in bottles covered with a wet yeasted molasses plate at 29 °C and the number of eggs was counted every 2 days (48 hours) for 10 days (modified from Beard et al., 2023). To assess the egg distribution in virgin conditions, ovaries were dissected after egg laying, and the location of eggs in the reproductive tract for each fly was examined.

For hatchability assays, one- to two-day-old virgin females were fed with standard food supplemented with wet yeast for a 2-day or 12-day feeding period (flipping to new food vials with wet yeast every four days) before the experiment. Once the feeding period ended, five virgin females were placed with ten *Oregon-R* males in bottles covered with a wet yeasted molasses plate at 29 °C and allowed to mate and lay eggs over 24 hours. Molasses plates with eggs laid for each condition were kept for an additional 24 hours at 25°C to allow eggs to hatch. The number of unhatched eggs per plate was quantified and subtracted from the total amount of eggs laid, allowing us to calculate the percentage of hatched eggs/plate.

### Quantification of mature follicles per ovariole

One-to-two-day-old virgin female flies with the desired genotype were fed with wet yeast paste for three or nine days, changing food every three days. At the end of the feeding period, ovaries were dissected in Grace’s medium and fixed in 4% EM-grade paraformaldehyde in PBT (1XPBS + 0.2% Triton X-100) for 15-20 min at room temperature. It is important to mention that during the dissections, ovary pairs were separated but with lateral oviducts attached to the base of the ovary to avoid loss of mature follicles from the ovarioles. The tissue was washed with PBT multiple times and stained with DAPI (final concentration 0.5ug/mL) for 15 minutes. Egg chambers were staged based on DAPI staining and morphology as previously described (Jia et al., 2016; Spradling, 1993).

### Statistical Analysis

Statistical tests were carried out using Prism 7 (GraphPad, San Diego, CA). Quantification results are displayed as mean ± SD or mean ± SEM as indicated in figure legends. Statistical analysis between two groups was conducted using a two-tailed Student**’**s t-test. Statistical analysis between more than two groups was conducted using One-way ANOVA. For the egg distribution assay used to calculate egg-laying time, Chi-square analysis was performed to assess significance.

## Acknowledgment

We would like to thank Drs. Andres Garelli, Alisson Gontijo, Maria Dominguez, Pierre Leopold, Dick Nässel for sharing fly lines and reagents; Bloomington Drosophila Stock Center and Vienna Drosophila Resource Center for fly stocks; and Developmental Studies Hybridoma Bank for antibodies. We thank Drs. Baosheng Zeng and Yuping Huang in the Sun laboratory for technical support. We appreciate constructive comments from anonymous reviewers. The Leica SP8 confocal microscope is supported by an NIH Award (S10OD016435) to Akiko Nishiyama. KY was a UConn Beckman Scholar. JS is supported by NIH/National Institute of Child Health and Human Development Grants (R01-HD086175 and R01-HD097206).

---

**Figure S1.**
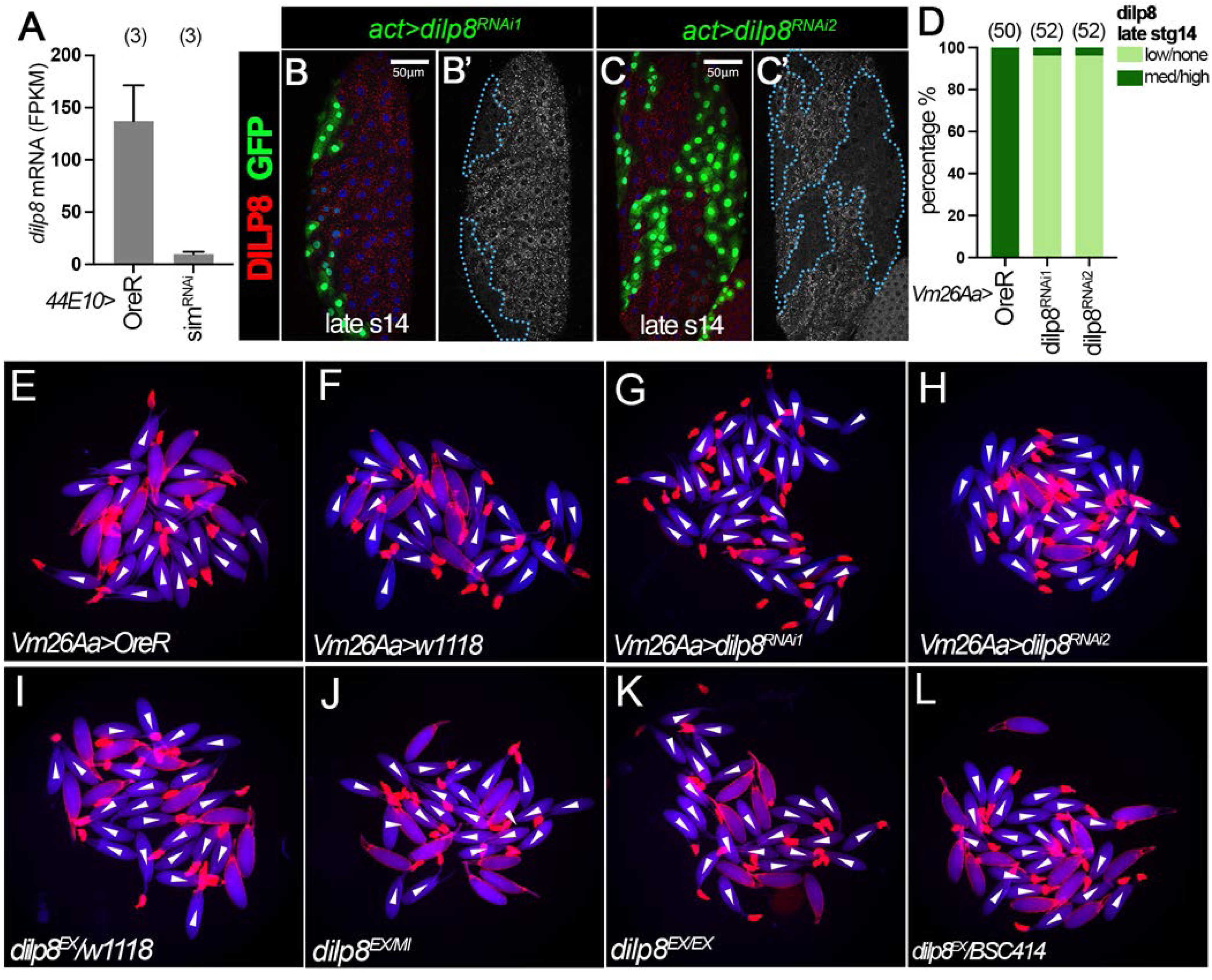
DILP8 can be efficiently depleted with follicle specific *dilp8^RNAi^* mediated knockdown. Flipout clones validate the DILP8A2 antibody specificity. Ex-vivo follicle rupture images display a lack of follicle rupture defects upon ovarian DILP8 depletion. (A) DILP8 mRNA expression levels (Fragments Per Kilobase of exon per Million mapped fragments, FPKM) from RNA Sequencing data collected with late stage-14 follicles from *OreR* control or *sim^RNAi^* females with *44E10-Gal4* and *Oamb-RFP.* The number of biological replicates is noted above each bar. **(B-C)** DILP8 protein expression (red in B-C and white in B’-C’) in late stage-14 follicles with flip-out clones (marked by GFP) overexpressing (B) *dilp8^RNAi1^* (*act>dilp8^RNAi1^*) and (C) *dilp8^RNAi2^* (*act>dilp8^RNAi2^*). The clone boundary is outlined by a blue dashed line. Nuclei are marked by DAPI in blue. The scale bar is 50um. **(D)** Quantification of DILP8 expression levels in late stage-14 follicles from *OreR* control, *dilp8^RNAi1^*, or *dilp8^RNAi2^* females with *Vm26Aa-Gal4* and *Oamb-RFP.* The number of follicles quantified is noted above each bar. **(E-L)** OA-induced follicle rupture results. (E-H) Representative images show mature follicles from *OreR* control (E), *w1118* control (F), *dilp8^RNAi1^*(G), and *dilp8^RNAi2^* (H) females with *Vm26Aa* and *Oamb-RFP* after 3-hour culture with 20 µM OA (OA+). (I-L) Representative images show mature follicles from *dilp8^EX^/w1118* heterozygous control (I)*, dilp8^EX/MI^* (J), *dilp8^EX/EX^* (K), or *dilp8^EX^/BSC414* (L) virgins after 3-hour culture with 20 µM OA (OA+). *Oamb-RFP* is shown in red; the bright-field signal is shown in blue. Ruptured follicles are labeled with white arrowheads and have corpus luteum accumulation at the posterior end by the dorsal appendage.

**Figure S2.**
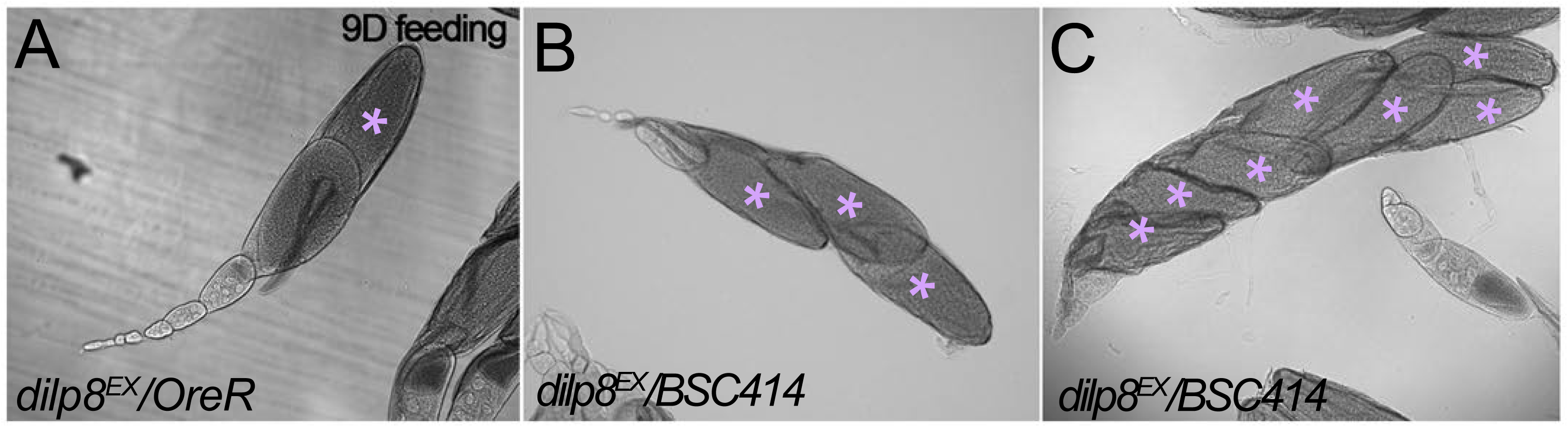
Mature follicle numbers increase in DILP8 depletion flies upon 9D feeding, even in the mated condition. (A-C) Representative brightfield images showing ovarioles from (A) *dilp8^EX^*/*OreR* heterozygous control or (B-C) *dilp8^EX^/BSC141* females after nine-day feeding with wet yeast (changing wet yeast and food every 3 days). Mature follicles are denoted with purple asterisks.

**Figure S3.**
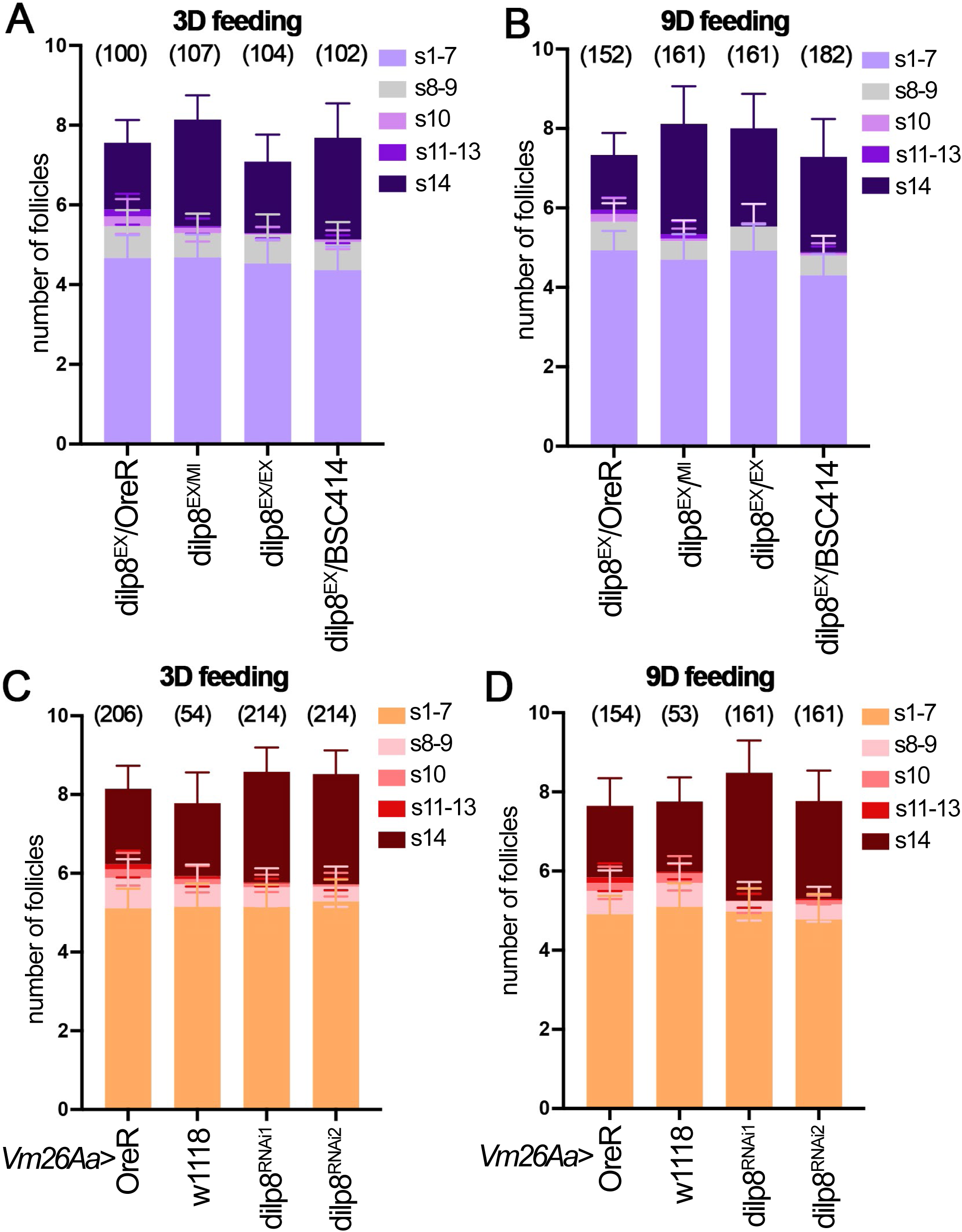
Ovariole follicle stage distribution in ovarian and global DILP8 depletion flies upon 3D or 9D feeding. (A-D) Follicle stage distribution per ovariole for virgins after 3-day (A, C) or 9-day (B, D) feeding. Dataset for *dilp8^EX^*/*OreR* heterozygous control*, dilp8^EX/MI^*, *dilp8^EX/EX^* or *dilp8^EX^/BSC141* mutant virgins (A-B). Datasets for *OreR* control, *w1118* control, *dilp8^RNAi1^*, or *dilp8^RNAi2^* virgins with *Vm26Aa-Gal4* and *Oamb-RFP* (C-D). The y-axis represents the percentage of ovarioles containing follicle stages within the following categories: stage 1-7, stage 8-9, stage 10, stage 11-13, or stage 14. The number of ovarioles quantified is noted above each bar. For all graphs, data points represent mean + SD.

**Figure S4.**
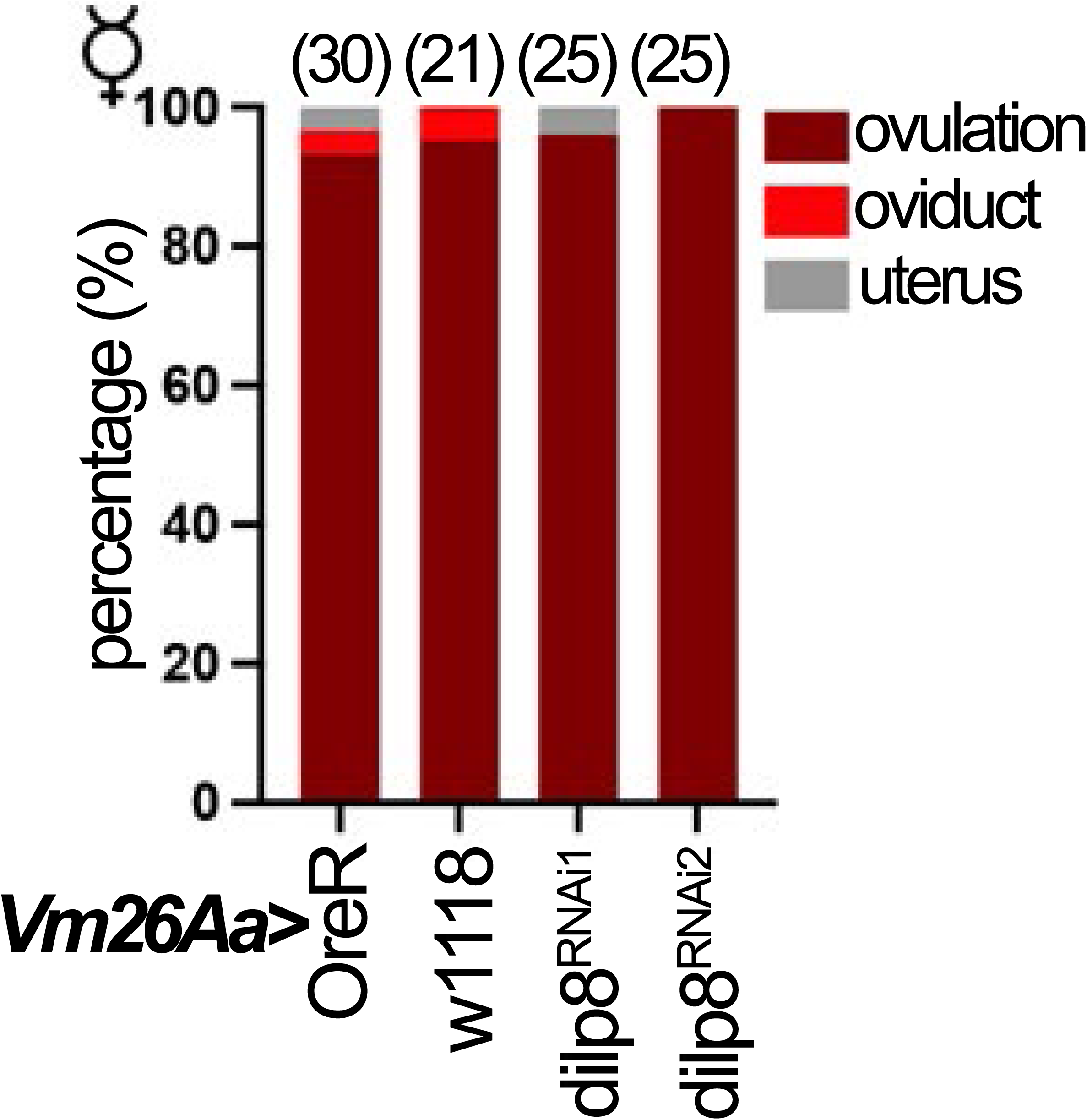
Eggs were present in the ovary at a higher incidence than other regions of the reproductive tract in DILP8-depleted virgin females relative to controls. Percentage of eggs located in the ovary, oviduct, and uterus across genotypes was assessed after the virgin egg-laying assay for *OreR* control, *w1118* control*, dilp8^RNAi1^*, or *dilp8^RNAi2^*virgin females with *Vm26Aa-Gal4* and *Oamb-RFP.* The number of females is depicted above each bar.

**Table S1.**
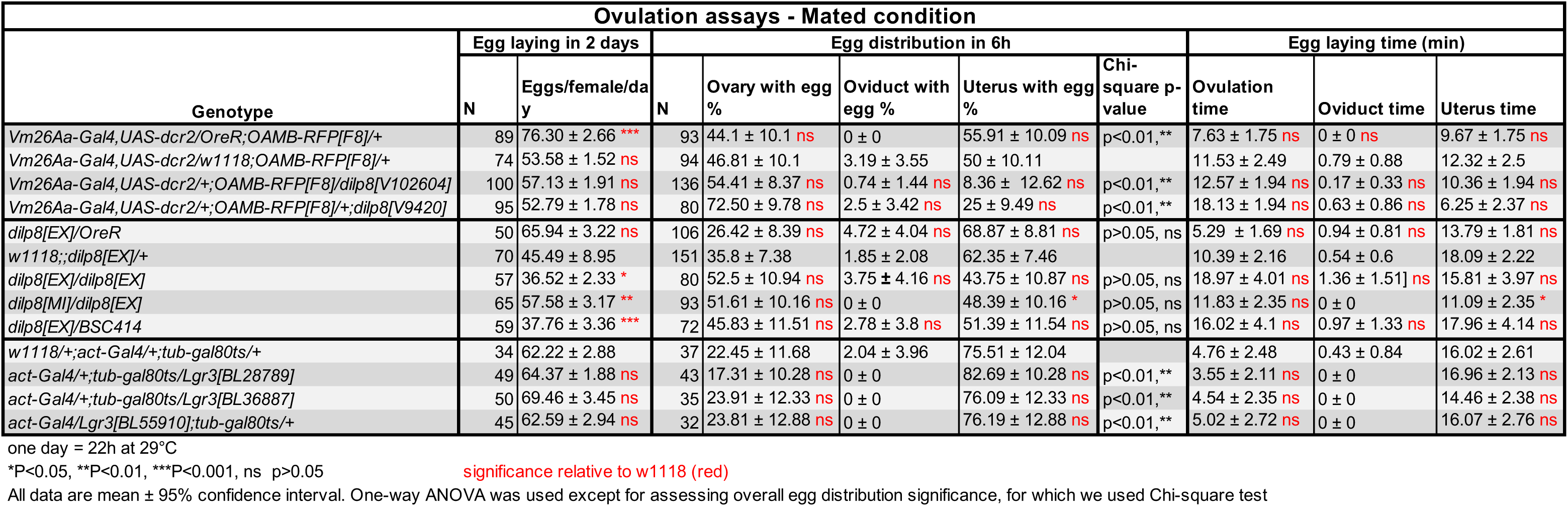
Egg laying, egg distribution in the reproductive tract, and egg laying time across multiple genetic conditions. Data are for Figure 1 and Figure 2.

